# Hierarchical Organization of Visual Feature Attention Control

**DOI:** 10.1101/2024.10.02.615879

**Authors:** Sreenivasan Meyyappan, Mingzhou Ding, George R Mangun

**Author notes:** **Corresponding Author:** George R. Mangun.

## Abstract

Attention can be deployed in anticipation of visual stimuli based on features such as their color or direction of motion. This anticipatory feature-based attention involves top-down neural control signals from the frontoparietal network that bias visual cortex to enhance the processing of attended information and suppress distraction. So, for example, anticipatory attention control can enable effective selection based on stimulus color while ignoring distracting information about stimulus motion. But as well, anticipatory attention can be focused more narrowly, for example, to select specific colors or motion directions that define task-relevant events and objects. One important question that remains open is whether anticipatory attention control first biases broad feature dimensions such as color versus motion before biasing the specific feature attributes (e.g., blue vs. green). To investigate this, we recorded EEG activity during a task where participants were cued to either attend to a color (blue or green) or a motion direction (up or down) on a trial-by-trial basis. Applying multivariate decoding approaches to the EEG alpha band (8-12 Hz) activity during the attention control period (cue-target interval), we observed significant decoding for both the attended dimensions (color vs. motion) <u>and</u> specific feature attributes (blue vs. green; up vs. down). Importantly, the temporal onset of the dimension-level biasing (color vs. motion) preceded that of the attribute-level biasing (e.g., blue vs. green). These findings demonstrate that the top-down control of feature-based attention proceeds in a hierarchical fashion, first biasing the broad feature dimension, and then narrowing to the specific feature attribute.

**Significance Statement:** During voluntary feature-based attention, electrophysiological and neuroimaging studies have highlighted the role of anticipatory (top-down) biasing of the sensory cortex in enhancing the selection of attended stimulus attributes, but little is known about how this is achieved. In particular, it is not clear whether attending to an attribute such as a color (blue vs. green) or motion direction (up vs. down) <u>first</u> biases all neural structures coding that dimension (color/motion) before biasing the specific attribute, or if the top-down signals directly bias only the attended attribute. Using EEG and multivariate decoding, we report that top-down attention control follows a hierarchical organization: first, the broader attended feature dimension is biased, which is followed by the biasing of the specific feature attribute.

## Introduction

Attention can be voluntarily deployed in anticipation of stimuli based on their location (spatial attention), features such as orientation, color, or motion direction (feature attention), and object properties such as faces, objects or landscapes (object attention). Functional imaging studies in humans have shown that in the period following an attention-directing cue but prior to the presentation of the target stimulus (cue-target interval or anticipatory period), the dorsal attention network (DAN), principally comprising of the intraparietal sulcus (IPS) and frontal eye fields (FEF), is activated. For non-spatial attention control such as attending to color or direction of motion of the stimuli, the inferior frontal junction (IFJ) is additionally activated, along with the DAN (Corbetta et al., 2000; Giesbrecht et al., 2003; Hopfinger et al., 2000; Kastner et al., 1999; Meyyappan et al., 2021; Nobre et al., 1997; Rajan et al., 2021; Szczepanski et al., 2013; Tamber-Rosenau et al., 2018). While studies have shown that the role of these regions is to issue top-down control signals to bias the sensory processing in favor of the attended information (Beauchamp et al., 1997; Bisley & Goldberg, 2010; Bressler et al., 2008; Chawla et al., 1999; Chen et al., 2012; Clark et al., 1997; Corbetta et al., 2008; Kastner & Pinsk, 2004; Martin et al., 1995; Nobre et al., 2006; O’Craven et al., 1997; Saenz et al., 2002; Schoenfeld et al., 2007; Sylvester et al., 2008; Treue & Martínez Trujillo, 1999; Wang et al., 2016), the temporal dynamics of how top-down control is achieved is not well established, in other words it is not clear whether anticipatory attention control first biases broad feature dimensions such as color versus motion before biasing the specific feature attributes (e.g., blue vs. green), or whether the specific to-be-attended feature attribute can be biased directly.

Insights can be gleaned from models of visual working memory. According to the dimensional feature bundle model (Töllner et al., 2015), the stimulus stored in working memory is represented by both the individual feature value and its feature dimension in a three-level hierarchy. In their model, the top level represents the stimulus identity, the second level represents the feature dimension such as color and shape, and the third level represents the specific feature attribute within each feature dimension. If, for example, the color of the remembered stimulus is to be retrieved from the working memory, then in addition to the activation of the representation of the specific color, the representations of the entire color dimension are also activated in the working memory (Niklaus et al., 2017). Specifically, the efficiency of reporting the probed stimulus was higher if it was within a dimension (color of object 1 to color object 2) than cross dimension (color of object 1 to orientation of object 2), suggesting that when a color is cued, all color sensitive units are enhanced, with additional enhancement for the cued or attended color, whereas the un-attended dimensions (orientation or shape or motion direction) are suppressed.

We hypothesize that a similar hierarchical organization exists for the voluntary control of feature selective attention. That is, when instructed to attend a specific attribute within a feature dimension (e.g., attend color blue, where color is the dimension and blue is the attribute), sensory enhancement of the feature dimension (e.g., color) will take place first (Stage-1), to be followed by sensory enhancement of the specific attribute (e.g., blue) (Stage-2). Other feature dimensions (e.g., motion) will be suppressed in Stage-1 while the other attributes within the attended dimension (e.g., yellow) will be suppressed in Stage-2. The alternative model is that sensory attributes such as color or motion direction may be represented independently in the attention control networks, and thus biased <u>directly</u> within visual cortex without following a hierarchical organization. In such a case, we would expect no temporal difference between the dimension level selection (motion or color) and the individual selection of motion direction (e.g., up vs. down) or color (e.g., blue vs. green).

To test the hierarchical organization model of feature attention control, we recorded electroencephalograms (EEG) from human participants performing a cued visual feature attention task. Each trial began with an auditory word cue that instructed participants to attend to one of two possible stimulus attributes (“blue” or “green”) within the color dimension, or to one of two possible attributes (“up” or “down”) within the motion dimension. Following a variable delay period, participants were presented with two intermingled streams of moving dots. One stream’s dots were moving upward and were either blue or green in color, and the other stream’s dots were in the other color moving downward. The dots in each moving stream could be either large or small in size, and the dot sizes in the two streams were independently manipulated. The participants’ task was to report the <u>size</u> of the dots in the stream having the attended color <u>or</u> motion direction while ignoring the other stream.

Following the prior literature demonstrating that EEG alpha oscillations (8 to 12 Hz) are modulated differentially depending on the features being attended (Snyder & Foxe, 2010), we used the temporal evolution of patterns of alpha oscillations to index the temporal dynamics of feature attention control. Specifically, we analyzed the alpha band activity using multivariate decoding approaches and generated the time courses of the decoding accuracy from decoding: (i) feature dimensions (color vs. motion), and (ii) decoding feature attributes (blue vs. green; upward vs. downward motion) within each feature dimension. By comparing the onset times of above-chance decoding we assessed whether the stage of biasing dimensions preceded that of biasing individual attributes within each dimension.

## Methods

### Participants

The study was approved by the Institutional Review Board of the University of California, Davis. Twenty-five right-handed participants (20 female; mean age: 24.83 years; range 19-39 years) with normal or corrected to normal vision and no known history of color blindness or neurological disorders participated in the study after providing written informed consent. This sample size was chosen using power analyses based on our previous published work (Nadra et al., 2023; Noah et al., 2020).

### Stimuli

The main stimuli involved two streams of centrally presented, spatially overlapping, and differently colored dots (blue or green) moving in opposite directions (up or down). In some experimental conditions, only one stream of dots appeared on the screen as described later (see next section). Within a stream (e.g., the up-moving stream), all the dots (n = 100) were of the same size (either large or small) and the same color (blue or green). However, across the two streams, the dots always had opposite colors and moved in opposite directions. For example, in a particular trial, if one stream of dots was <u>green</u> and moved <u>upwards</u>, then the other stream of dots would be <u>blue</u> and moved <u>downwards</u>. The sizes of the dots across the two streams were pseudo-randomly determined and <u>independent</u> of each other. In other words, in a trial, if the upward moving dots were <u>large</u>, then the dots in the downward moving stream could either be large <u>or</u> small. Given that the task was to discriminate if the dots in the attended stream were large or small (see below), this design ensured that the participants had to attend the cued stream to perform the task.

### Experimental Paradigm

Participants performed a cued feature-attention task of fifteen runs of 3 mins each in an electrically shielded, sound-attenuating chamber (ETS-Lindgren). As shown in Figure 1, a fixation point was placed at the center of the monitor, and the participants were instructed to maintain fixation on the fixation dot during the experiment. At the beginning of each trial, one of the four auditory cues was presented (attend motion cues: spoken words “up” or “down;” attend color cues: spoken words “blue” or “green”), instructing the participants to direct attention to a stream of moving dots based either on their direction of motion (up or down) or color (blue or green). From trial to trial, the cues were pseudo-randomly determined such that the number of trials per condition were identical but their order of presentation in a block was fully randomized. The mean duration of the four auditory cues was 530 ± 34 ms. Targets appeared after a random delay (cue-target onset asynchrony interval) of 1250 to 1750 ms (time-locked to cue-onset). On 80% of the trials, referred to as the *<u>dual-stream</u>* condition, two streams of oppositely moving and oppositely colored dots appeared for 250 ms. The participants were instructed to discriminate the <u>size of the dots</u> in the <u>cued stream</u> (target) while ignoring the size of dots in the un-cued stream (distractor), and to press a button to indicate whether the dots in the cued stream were large or small.

While the <u>dual-stream</u> condition occurred 80% of the time, for the remaining 20% of trials, the stimuli consisted of only <u>one</u> stream of dots referred to as the *<u>mono-stream</u>* condition. In these trials, the dots were either <u>only</u> in the cued color or motion direction (*<u>mono-valid</u>; 10*% of trials) with the distractor stream absent <u>OR</u> in the un-cued color/motion direction (*<u>mono-invalid</u>;* 10% of trials) with the target stream absent. The task during the <u>mono-valid</u> trials stayed the same as the dual-stream trials, namely, the participants had to discriminate the size of the dots. For *<u>mono-invalid</u>* trials, when the cued stream was not present, participants were instructed to <u>shift</u> their attention to the <u>un-cued stream</u> and report the size of the dots. The mono-stream conditions (valid and invalid) were included to obtain behavioral measures of the benefits of the attentional cueing. Analogous to the behavioral effect of cue validity observed in the standard attention cueing paradigms (Posner et al., 1980), we predicted that participants would be slower to respond on invalidly cued mono-stream trials. A pseudo-randomly distributed intertrial interval (ITI) between 2000 and 3000 ms separated target offset from the cue onset of the next trial. A training session preceded the main study to familiarize the participants with the task.

### EEG Recording

Continuous EEG data was acquired with a 64-channel Brain Products actiCAP snap active electrode system (Brain Products GmBH) and digitized using a Neuroscan SynAmps2 input board and amplifier (Compumedics USA Inc.). Signals were recorded with Curry 8 acquisition software with a sampling rate of 1000 Hz and a bandpass filtering between 0 to 400 HZ online. Water soluble electrolyte gel was used to maintain surface contact between the electrode and scalp; the electrode impedances were maintained at less than 25 KΩ.

**Figure 1:**
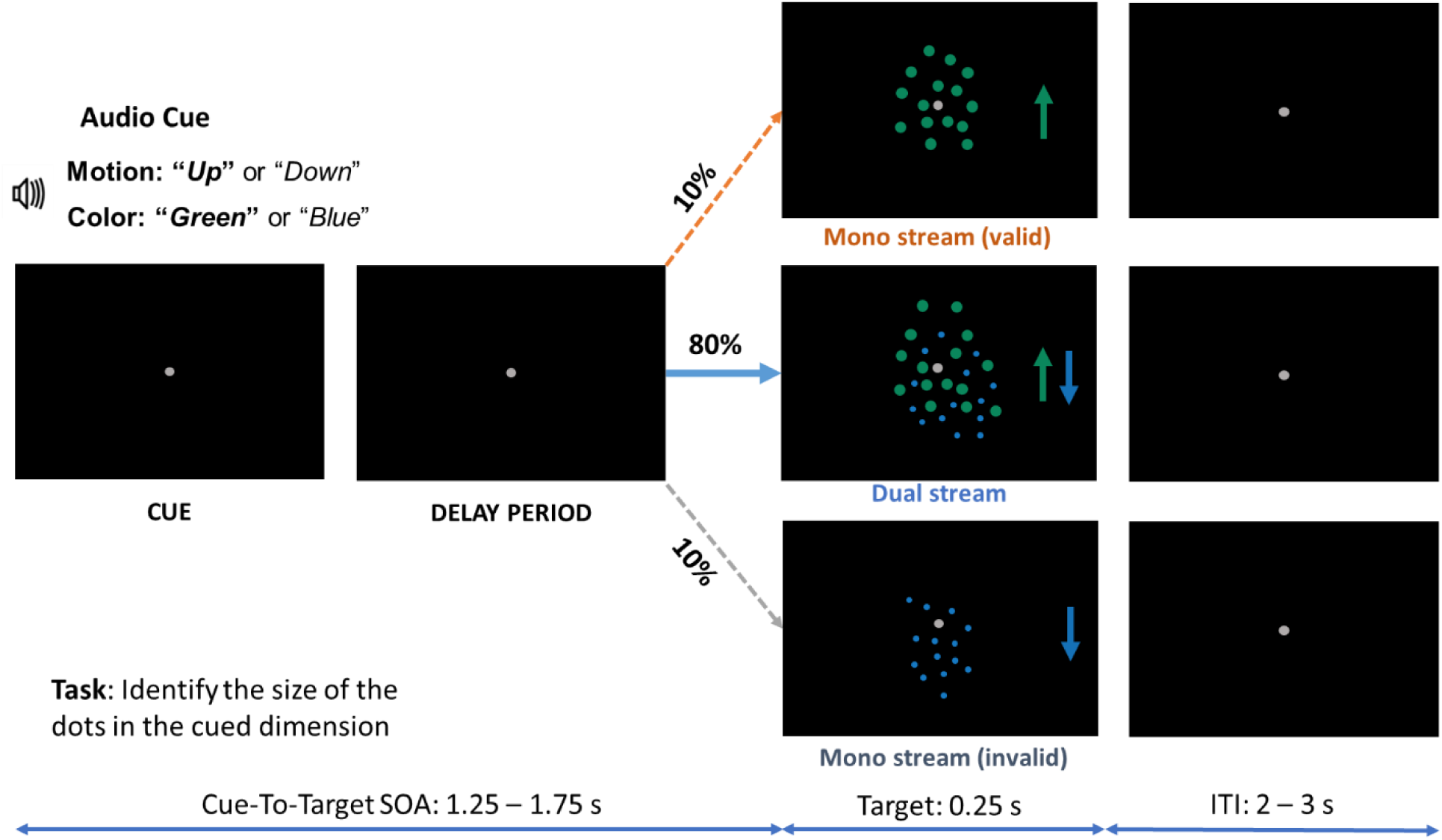
Experimental paradigm. An auditory cue directed participants to selectively and covertly attend either a color or a motion direction in order to detect the size of the dots in the attended stream. On 80% of trials, this was followed by a compound color-motion stimulus, such that task performance relies on using the cue information; distractor presence (the un-cued attribute) provided competition that requires attention. On 20% of trials, only the cued (valid) or un-cued (invalid) stimuli were presented, to provide behavioral measures of attention cueing (valid vs invalid). The different cue-target types are randomly interleaved <u>within</u> blocks. In the example given above, if the cue was “*Up*” or “*Green,*” the correct button press in valid trials would be to indicate the dots being <u>large</u>.

### EEG Preprocessing

All data preprocessing procedures were completed with the EEGLAB toolbox written in MATLAB (Delorme & Makeig, 2004). For each participant, the EEG data files from the individual runs were merged into a single dataset before data preprocessing. The data were Hamming window sinc FIR (finite impulse response) filtered (1–83 Hz) with -6 dB roll off at 0.5 and 93 Hz, and then down-sampled to 250 Hz. Data were re-referenced to the average of all electrodes (common average reference) and epoched from 1000 ms before cue onset to 4000 ms after cue onset to include both cue and stimulus-evoked neural data. An Independent component analysis (ICA) routine implemented in the EEGLAB was used to remove blink, muscular and oculomotor artifacts.

A current source density (CSD) transform was applied to the epoched data by estimating the surface Laplacian to reduce the influence of volume conduction and common reference from the EEG data (Tenke & Kayser, 2012). To extract alpha band activity (8-12 Hz) as a function of time, the epoched CSD data was bandpass filtered between 8-12 Hz and Hilbert-transformed to obtain the alpha-band complex analytic signal, whose magnitude was then squared to yield the alpha power time course.

### Decoding Analysis

The alpha power time course from the epoched CSD data was used for the decoding analysis. To reduce noise, a moving average filter was applied, in which the epoched EEG (CSD) data was temporally smoothed by computing the average of five adjacent timepoints (20 ms time window). The decoding was done on the smoothed data at every time point. The 64 alpha power values from sixty-four channels were used as 64 input features to the classifiers. A linear support vector machine (SVM) was applied to classify the attention conditions by using the trial-averaged decoding approach (Bae & Luck, 2018). For decoding two attention conditions (i.e., attend-up vs. attend-down; attend-blue vs. attend-green; attend-motion vs. attend-color), the data within each attention condition were divided into three parts (three-fold cross-validation). The trials in each part were averaged for a given timepoint to boost the signal-to-noise ratio (Bae & Luck, 2018). The classifier was trained using the averaged alpha power from two parts of each attention condition and tested on the remaining part. To avoid bias in grouping of the trials into training and testing set, we repeated the process of dividing the trials into two-part training and one-part testing sets twenty-five times. The decoding performance was measured in terms of average decoding accuracy across the twenty-five repetitions. Above chance decoding accuracy is taken as evidence that the two attention control states differed in the pattern of alpha, reflecting the differential biasing of visual cortex for the two contrasted attention conditions. The higher the decoding accuracy, the more distinct the patterns of alpha power distribution, the more distinct the two attention control states.

To determine whether the decoding accuracy at a particular timepoint is above the chance level we compared the actual decoding accuracy against the chance level accuracy (50%) using paired t-tests. The decoding was deemed significantly above chance level if it was higher than the cluster-corrected threshold determined by requiring p<0.05 (Fahrenfort et al., 2017). To avoid any potential bias exerted by the unequal numbers of trials in different types of comparisons (e.g., comparing attend-blue vs. attend-green with attend-color vs. attend-motion), we <u>subsampled</u> and randomly chose 50% of the trials from motion (up and down combined) and color (blue and green combined) conditions for motion versus color decoding analysis. The decoding analysis with subsampling was repeated 25 times for each participant and the resulting 25 decoding accuracies were averaged.

### Estimating Decoding Onset Latencies

A jack-knife approach (Miller et al., 1998) was implemented to estimate the time at which decoding accuracy rises above chance level. The following steps were performed to estimate the decoding onset latency: (1) Decoding accuracy time course from a subsample of N-1 (N = 25) participants was extracted by excluding one participant. For example, in the first iteration, the decoding accuracy time course from subject 1 was excluded and the decoding accuracy time courses from subjects 2 to 25 (N = 24) were considered. (2) The distribution of the decoding accuracy time course from the subsample were compared against a chance-level and significant decoding clusters (p<0.05) were obtained. (3) Decoding clusters earlier than 50 ms after the cue onset were excluded from the analysis since because top-down feature attention control is known to take longer than 50 ms after the onset of the cue to begin being implemented (Grent-’t-Jong & Woldorff, 2007; Liu et al., 2007; Nakayama & Mackeben, 1989). (4) Onset latency was determined by noting the <u>first</u> timepoint of the <u>earliest</u> significant cluster. (5) Steps (1-4) were repeated 25 times—by leaving out a subject exactly once—to obtain 25 decoding onset latencies. The decoding onset latency distribution was compared across different comparisons.

### Eye-tracking Data Recording and Preprocessing

An SR Research EyeLink 1000 eye tracker system was used to record the eye-movements and pupillary activity at a sampling rate of 1000 Hz. During pre-processing, eye-blinks were detected and gaze-positions during eye blinks were determined by a cubic spline interpolation algorithm. The continuous eye-movement data (x and y coordinates) was epoched from 200 ms before cue onset to 1200 ms after cue onset (-200 ms to 1200 ms).

## Results

### Behavioral Analysis

In order to establish that the subjects were using the cue information to selectively bias attention, we analyzed their performance during the <u>mono stream</u> trials, that were 20% of the total trials. For <u>validly cued</u> mono-stream trials <u>(distractor absent)</u> (10%), the average response times (RT) were 779.3 ± 135.4 ms for motion trials and 758.1 ± 122.9 ms for color trials. The response accuracy for motion and color trials was 89.8 ± 7.4% and 88.7 ± 9.0%, respectively. Performance in validly cued trials did not differ significantly between attended feature dimensions (motion vs. color) (see below for the main effects of attention type). For <u>invalid mono-stream trials (cued target absent)</u> (10%), where the participants had to shift attention to the un-cued stream and report the size of the dots, the response times (RT) were 892.5 ± 188.2 ms for motion trials and 899.5 ± 188.7 ms for color trials. The response accuracy for motion and color trials was 83.1 ± 13.0% and 85.5 ± 14.1%, respectively. Performance on invalidly cued trials did not differ significantly between the two attended feature dimensions (motion vs. color) (see below for the main effects of attention type).

To assess the effect of cue validity, we compared validly cued mono-stream trials and invalidly cued mono-stream trials using two-way ANOVAs: cue validity (valid vs. invalid) by attention type (motion vs. color). For reaction time (RT), we found a statistically significant main effect of cue validity (valid vs. invalid; Figure 2A) [*F_validity_* _(1,24)_ = 15.52, *p* = 2 × 10^−4^, η^2^ (effect size) = 0.14]. There were no statistically significant main effects of attention types (motion vs. color) for RTs (*F_attention type_* _(1,24)_ = 0.05, *p* = 0.83, η^2^ = 3.42 × 10^−4^), nor were there significant interactions between cue validity and attention type (*F_validity × attention type_*_(1,24)_ = 0.19, *p* = 0.66, η^2^ = 1.71 × 10^−3^). For accuracy, we found a statistically significant main effect of cue validity (valid vs. invalid; Figure 2B), with validly cued trials having higher accuracy than invalidly cued trials [*F_validity_*_(1,24)_ = 4.85, *p* = 0.03, η^2^ = 0.046]. There were no statistically significant main effects of attention types (motion vs. color) on accuracy (*F_attention type_* _(1,24)_ = 0.08, *p* = 0.77, η^2^ = 7.81 × 10^−4^), nor were there significant interactions between cue validity and attention type (*F_validity × attention type_* _(1,24)_ = 0.61, *p* = 0.43, η^2^ = 5.46 × 10^−3^).

**Figure 2.**
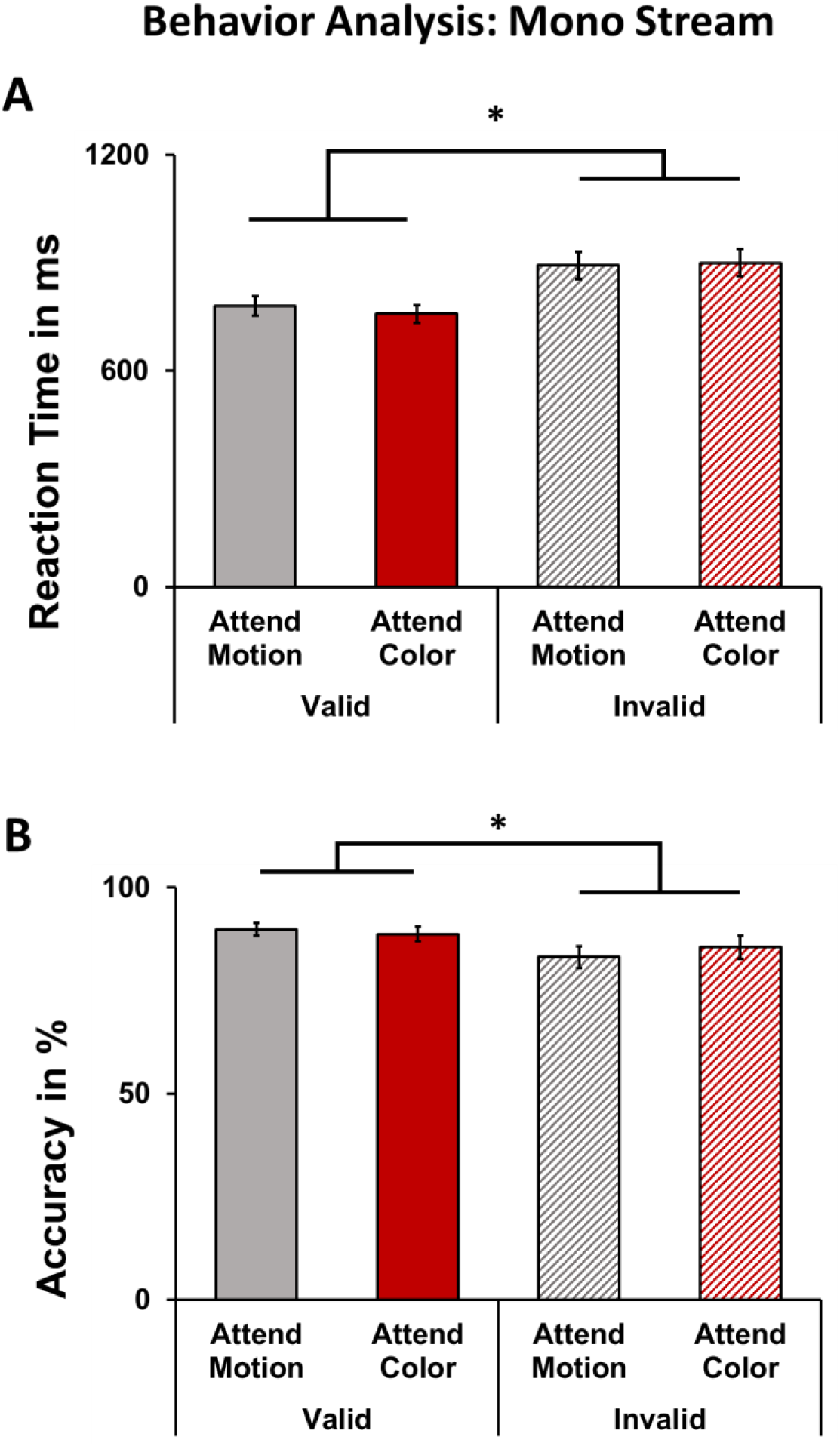
Behavioral analysis: mono-stream trials. (A) RTs to the targets (collapsed over motion direction and color) were significantly faster when validly cued targets were compared to invalidly cued targets. (B) Accuracy (% correct) was significantly higher for validly cued trials compared to invalidly cued trials. The error bars denote SEM and *p<0.05.

*<u>Dual stream:</u>* Eighty percent of the trials in the experiment were dual-stream trials where both cued and un-cued stimulus streams were present, and participants attended to the cued stream (target) while ignoring the un-cued stream (distractor). To assess whether there were any differences in arousal or task-difficulty among the four different attention trial types (attend-up, attend-down, attend-blue, and attend-green), we conducted one-way ANOVA with four levels (the four attention conditions) for RT and accuracy measures separately(Figure 3). Neither RT nor accuracy was found to be significantly different among the four attention trial types: RT = [F*_RT_*_(3,96)_ = 0.87, *p* = 0.45, η^2^ = 0.02]; accuracy = [F*_accuracy_* _(3,96)_ = 1.3, *p* = 0.28, η^2^ = 0.04]. These patterns for RT and accuracy suggest that the attention conditions did not differ in overall task difficulty or arousal across different attention conditions. Taken together, these behavioral analysis results suggest that the participants deployed covert attention to different feature attributes based on the information in the cues.

### Alpha Topography Analysis

For cue-evoked neural activity, we first examined the distribution of alpha power as a function of time over the scalp for different attention conditions. A pairwise difference map in alpha power was then computed using the CSD transformed and epoched EEG data for different feature attention conditions, and the averaged group-level difference maps were visualized in topographic head plots for different time periods in the cue-target interval. In Figure 4A, comparing the scalp topography of attend-motion against attend-color (subtracting the alpha power topography of one condition from that of the other), beginning in the 400-600 msec interval, we observed a decreased alpha power (blue colors) for attend motion in the parietal channels that continued to develop bilaterally over time during the cue-target interval for several hundred millisecond. This decrease in alpha in the dorsal parietal channels for attend-motion relative to attend-color towards the later stages of the cue-target interval aligns with the findings of Snyder and Foxe (2010) who also reported differential sources in the dorsal and ventral stream for the modulation of alpha power following attend-motion and attend-color cues, respectively. Additionally, comparing attend-up versus attend-down (Figure 4B) or attend-blue versus attend-green (Figure 4C) also yielded different alpha power patterns, suggesting that alpha power distribution over the scalp with the to-be-attended feature attributes within an attended feature dimension (up vs. down in the motion dimension or blue vs. green in the color dimension). To quantify these effects more precisely with higher temporal resolution, we turned to the MVPA decoding approach using alpha power from 64 scalp electrodes as input features, thus yielding decoding accuracy time courses for build of alpha power differences as a function of attention condition.

**Figure 3.**
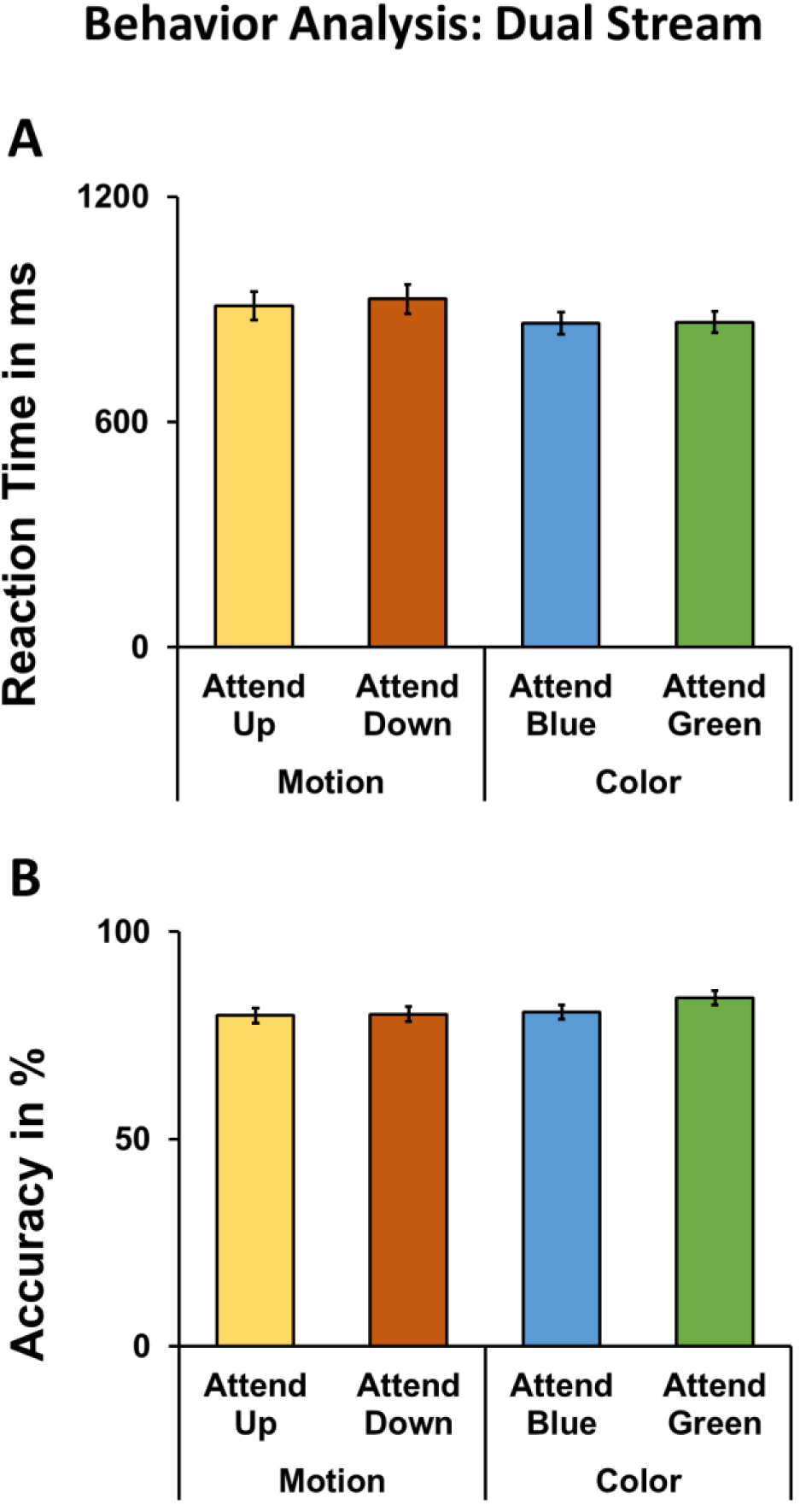
Behavioral analysis: dual stream trials (when both the target and distractor dot steams were present). Both reaction times and accuracy were similar across attention conditions (attend-up, attend-down, attend-blue, and attend-green). Error bars denote SEM.

### Decoding the Attended Feature Dimensions

Cue-evoked alpha patterns were used to distinguish the attended feature dimensions (motion vs. color). The time course of decoding accuracy is shown in Figure 5A. The decoding accuracy was at chance level at the start of the cue-target interval and rose above chance level at 180 ms after cue onset, and barring a brief period of 75 ms (between 275 ms – 350 ms), the decoding accuracy remained significantly above chance level for the analyzed cue-target period (0 to 1200 ms). This period was chosen to allow all trials including the trials with the shortest cue-target interval of 1250 ms to be included in the analysis.

### Decoding the Attended Feature Attributes Within an Attended Dimension

To investigate whether the modulations of alpha oscillations encode specific attended feature attributes within an attended feature dimension, we next trained linear SVM models to classify attend-up versus attend-down within the motion dimension or attend-blue versus attend-green within the color dimension. The average time courses of decoding accuracies for attend-up versus attend-down and attend-blue versus attend-green are shown in Figure 5B. The earliest time point that decoding accuracy for feature attributes reached significance was ∼384 ms after cue onset.

**Figure 4.**
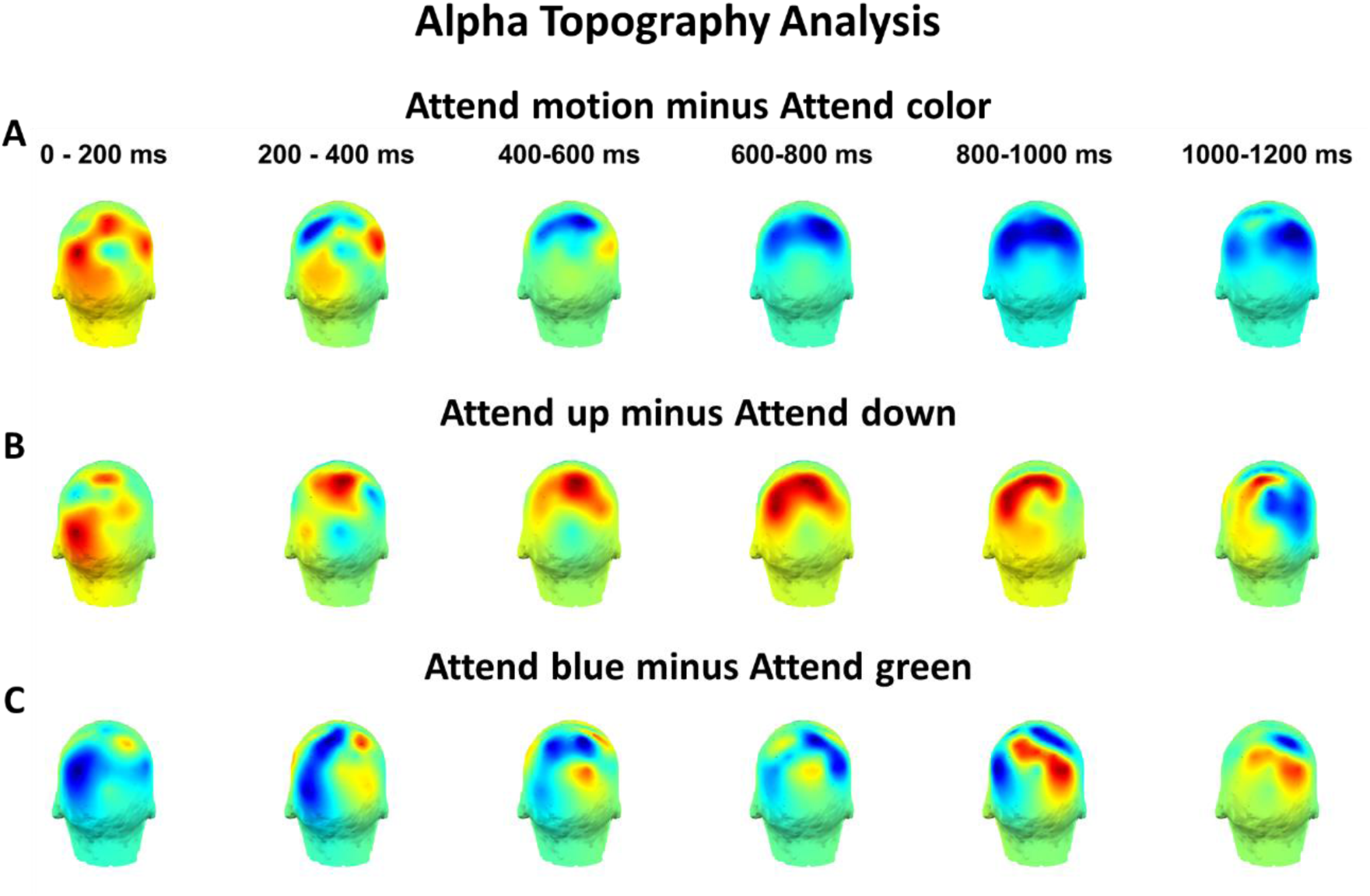
Alpha power difference at different time points displayed as topographic maps. The maps are views from behind the head, thus focusing on parietal and occipital scalp regions. (A) Attend-motion minus attend-color. (B) Attend-up minus attend-down. (C) Attend-blue minus attend-green. Here time 0 denotes the onset of the attention-directing cue. Note: The mean duration of the auditory cues was 530 ± 34 ms, and thus the patterns in the maps developing after 400-600 ms represent the post-cue/pre-target anticipatory period.

**Figure 5.**
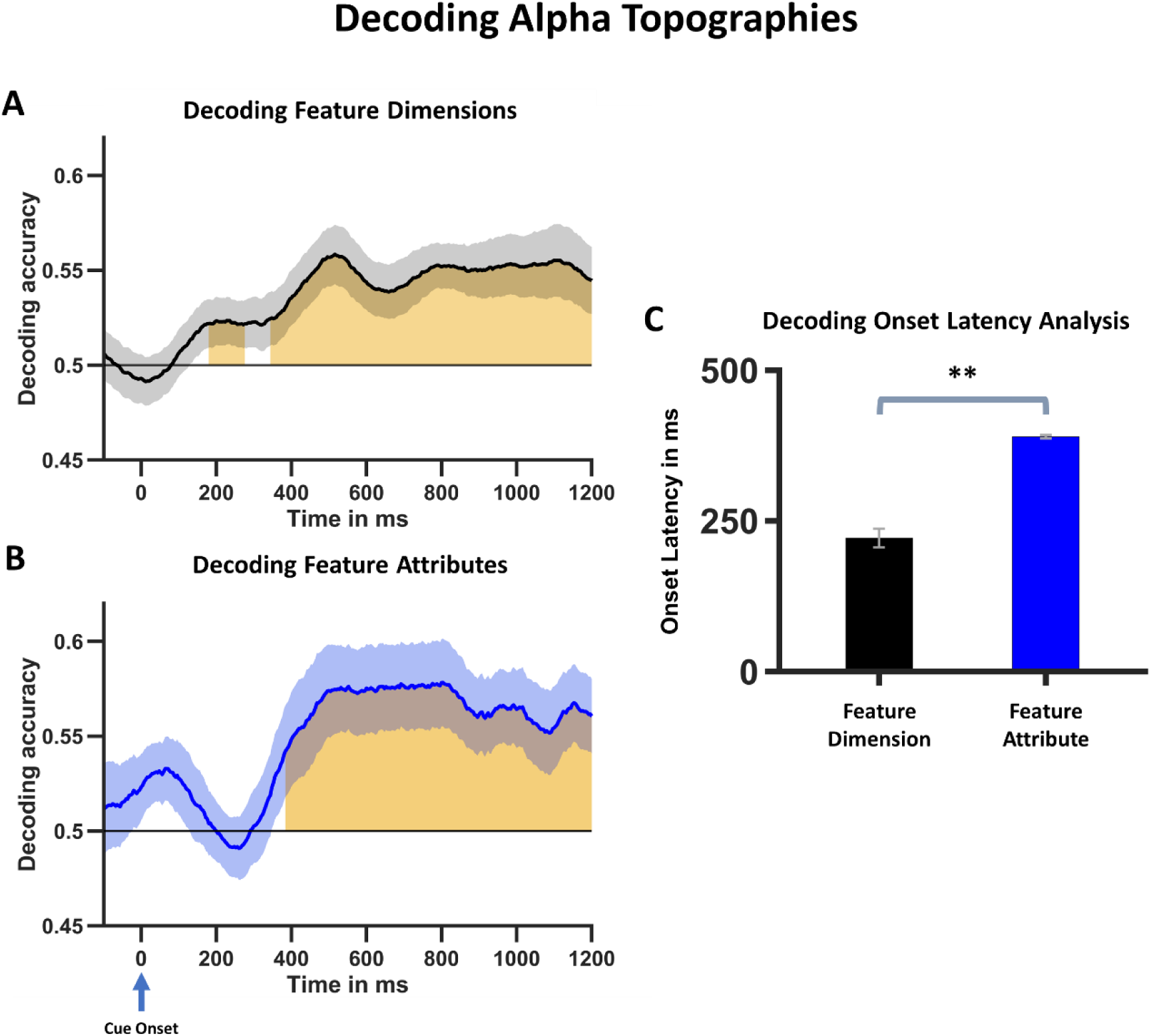
Decoding alpha topographical patterns during the cue-target interval. The accuracy time courses from decoding attended dimensions (attend-motion vs. attend-color; panel A) and decoding attended feature attributes which is the average of attend-up vs. attend-down and attend-blue vs attend-green decoding timecourses (panel B) with onset of cue at 0 sec. The chance level accuracy was 0.5. The shaded area denotes time points where the classifier performance was above chance level (p<0.05 cluster corrected). (C) The decoding onset latencies for decoding attend-motion vs attend-color (dimension decoding) were significantly earlier compared to decoding the attended feature attributes. Here time 0 denotes the onset of the attention-directing cue. The error bars and shaded regions around the decoding timecourses denote SEM and **p<0.0001.

### Comparison of Decoding Onset Latencies

In the decoding approach, when the decoding accuracy between two attention conditions rose and remained above chance level for the first time, referred to here as the decoding onset latency, it signifies the beginning of the formation of distinct neural representations of the two attention control states. Inspection of Figure 5A and 5B suggests that the formation of the neural representations of the attended feature dimensions (motion vs. color) and that of the specific attended feature attributes within an attended dimension started at different times. As shown in Figure 5C, the onset latency of decoding attend-motion vs attend-color significantly (p<0.0001) <u>preceded</u> the onset latency of decoding of the attended feature attributes (attend-up versus attend-down and attend-blue versus attend-green). Because attend-up and attend-down trials were combined as attend-motion trials and attention-blue and attend-green trials were combined as attention-color trials, there is a possible concern that the latency effect in Figure 5C could have been driven by different numbers of trials used in different analysis. We mitigated this concern by performing decoding analysis for attend-motion versus attend-color (dimension classification) using only 50% of the trials (random subsampling), thereby equalizing the numbers of trials used for decoding attended dimensions and that used for decoding attended attributes within an attended dimension. Such subsampling was performed 25 times and the 25 subsampled decoding accuracies were averaged to obtain the attend-motion versus attend-color decoding timecourse.

**Figure 6.**
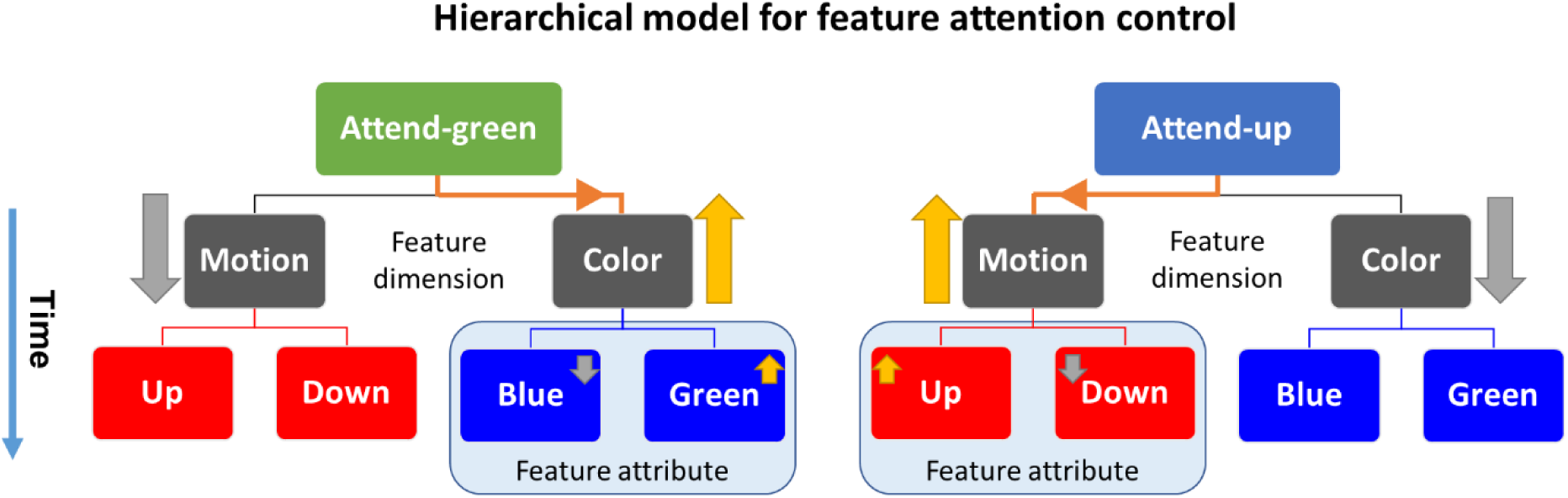
Schematic representation of the hierarchical organization of feature attention control where the selection of feature dimensions temporally preceded the selection of feature attributes. The orange arrow indicates the example top-down control scenarios (left: attend green; right attend up). The yellow and gray arrows indicate the facilitated and suppressed dimension (stage I) and attribute (stage II).

These latency data suggest a hierarchical organization of feature attention control which consists of two stages (Figure 6). In the first stage, the broad attended category is biased such that the attended feature dimension is relatively facilitated and the unattended (ignored) feature dimension is suppressed. In the second (later) stage, the biasing narrows to the specific to-be-attended motion direction (up vs. down) or color (blue vs. green) and suppresses the unattended direction or color.

### Patterns of eye movements

Previous work has shown that eye movement patterns can affect decoding analysis from EEG (Hong et al., 2020), MEG (Cichy et al., 2015) and fMRI (Thielen et al., 2019) data. To examine whether there were systematic eye movement patterns that distinguished different attention conditions, we analyzed the eye tracking data using linear SVMs (Meyyappan et al., 2021; Rajan et al., 2021). The decoding accuracy was at chance level in the cue-target interval throughout the 0 to 1200 ms period for all pairs of attention conditions: attend motion versus attend-color (Figure 7A), attend-up versus attend-down (Figure 7B), and attend-blue versus attend-green (Figure 7C). These eye-tracking results rule out the possibility of systematic eye movements influencing the alpha-decoding results.

**Figure 7.**
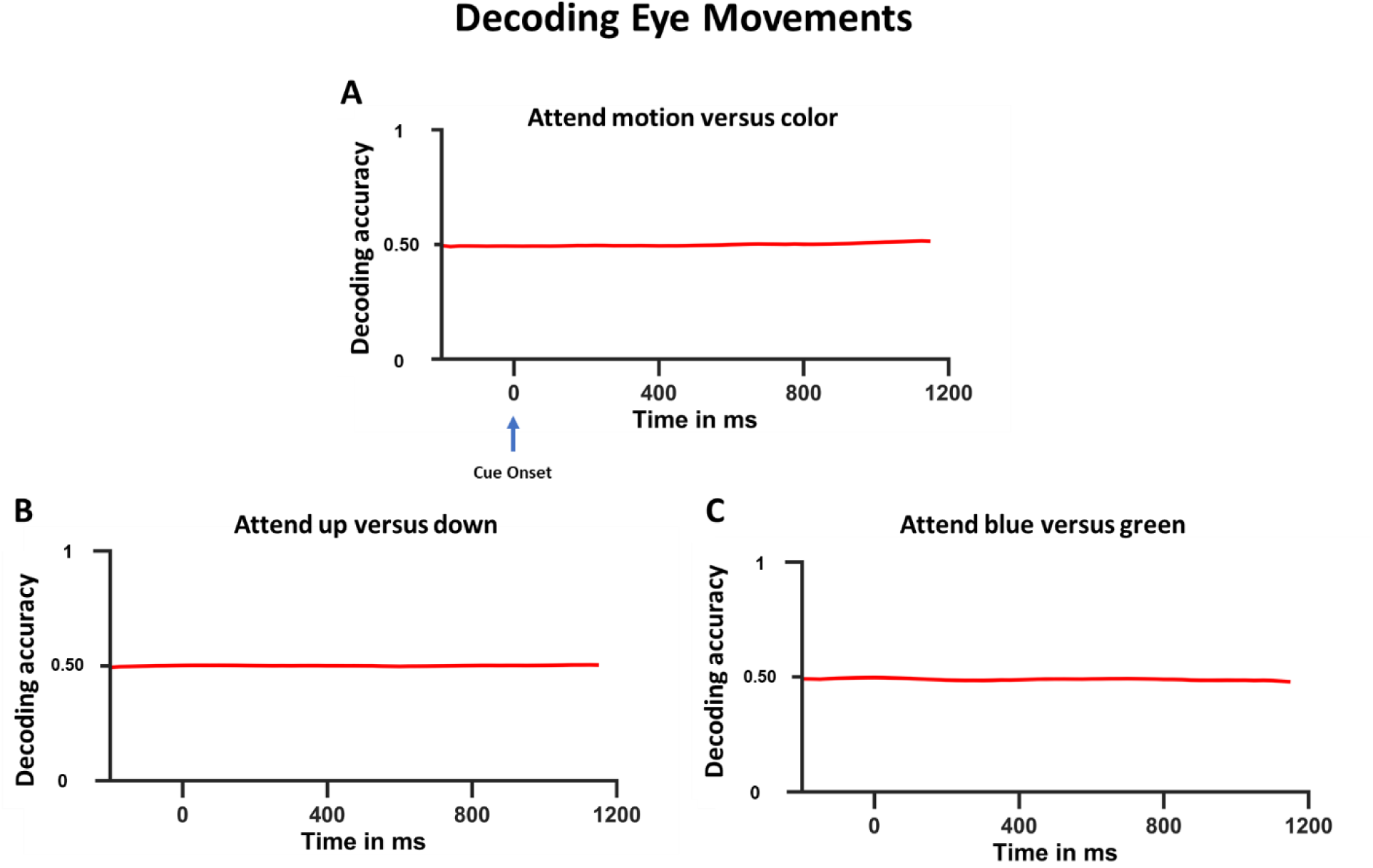
Decoding eye movements. (A) Decoding accuracy as a function of time for attend motion versus attend color (feature dimension classification). (B) Decoding accuracy as a function of time for attend up versus attend down. (C) Decoding accuracy as a function of time for attend blue versus attend green. Here, time 0 refers to the onset of the cue and the decoding accuracies were not above chance level throughout the cue-target interval for all three classifications.

## Discussion

Using spatial patterns of alpha oscillations (8 to 12 Hz) as neural observables, we applied multivariate decoding approaches to investigate if there is a hierarchical organization underlying feature attention control. First, comparing the univariate alpha scalp topographies, we found different patterns of EEG alpha topographies for attending different feature dimensions (i.e., attend-motion vs. attend-color) as well as for attending different attributes within a feature dimension (attend-up vs. attend-down within motion and attend-blue vs. attend-green within color) in the cue-target interval, demonstrating that patterns of alpha oscillations contain information on different attention control states. Second, comparing the time courses of the decoding accuracy, we found that, depending on the comparison, above-chance decoding starts at different times relative to the cue onset: attend-motion versus attend-color (attend different feature dimensions) begins at ∼180 ms, while attention to specific feature attributes begins at ∼385 ms. This latency data is taken to suggest that feature attention control is hierarchically organized: it was first deployed to the attended feature dimension by enhancing the attended feature dimension and suppressing the unattended feature dimension, and then to the attended feature attribute within the attended feature dimension by enhancing the attended feature attribute and suppressing the unattended feature attribute.

### Patterns of alpha oscillations as indices of feature attention control

The modulation of alpha oscillations by visual spatial attention is well-established (Popov et al., 2019; Rihs et al., 2007; Samaha et al., 2016; Sauseng et al., 2005; Thut et al., 2006; Worden et al., 2000). During visual spatial attention control, alpha power is suppressed over the posterior cortex corresponding to the attended location, reflecting increased readiness for processing the impending visual input (Fiebelkorn & Kastner, 2020; Popov et al., 2017). Increased alpha, on the other hand, is thought to be a neural mechanism for suppressing the cortical regions not relevant for the processing of the impending visual input. Snyder and Foxe (2010) were among the first to report the involvement of alpha oscillations during non-spatial (feature) attention. In a cue-target paradigm similar to the one used here, participants were cued to covertly attend either motion or color on a trial-by-trial basis, and upon the presentation of colored moving dots, reported if any deviations occurred in the attended feature dimension (color or motion). Extracting color and motion-sensitive alpha components from the cue-target interval, they identified the putative neural sources for the increased alpha power within the ventral or the dorsal visual pathways when color or motion was to be suppressed, in line with the Gating by Inhibition Model as the role of alpha oscillations in facilitating top-down feature-based-attention (Jensen & Mazaheri, 2010). Our univariate analysis comparing the alpha activity between attend-motion (attend-up and attend-down combined) and attend-color (attend-blue and attend-green combined) revealed different topographic patterns of alpha power modulation, depending on the feature dimension being attended, consistent with the findings of Snyder and Foxe (2010); our multivariate decoding of feature dimensions is also consistent with those findings. In addition, going beyond the work of Snyder and Fox, we additionally found that the topographic patterns of alpha oscillations distinguished specific attended feature attributes within an attended feature dimension, and these patterns could be decoded using multivariate methods.

We note that this finding is in contrast with the recent findings of Gundlach et al. (2023) who report no attribute (color) specific modulation in alpha oscillations. Gundlach and colleagues presented four sets of moving dot patterns with two different colors (blue and orange) where one set of blue and orange dots was presented at the center of the screen and the remaining blue and orange set were presented at the left or right of the screen. The dot patterns were present throughout the experiment and were set to flicker at different frequencies. At the start of a trial, following a variable pre-cue period, participants were cued to attend to a color (blue or orange) in the centrally presented dots and detect when a coherent motion where 40% of dots briefly moved (300 ms) in one common direction. Analyzing the cue evoked SSVEP amplitudes and alpha band activity as a function of the attended color, Gundlach et al. showed that SSVEP amplitudes were significantly greater for the attended color than un-attended color both for the centrally presented stimuli and peripherally presented stimuli. However, for the alpha-band activity, no significant differences were observed in alpha desynchronizations for the attended and the unattended color in their contralateral electrodes. Our findings, on the contrary, suggest that alpha band activity encodes the attended feature attribute during the anticipatory period. We believe two key differences between our findings and those of Gundlach et al. may account for the differential findings. The first difference concerns univariate versus multivariate approaches.

Gundlach and colleagues used a univariate approach to index EEG alpha activity; univariate scalp topographies, while an informative tool, do not lend easily to quantitative analysis and are less sensitive to subtle differences at a pattern level. The multivariate approach, which combines alpha power topography with machine learning-based classification, overcomes these limitations and may enable insights not possible with univariate methods. Second, the visual stimuli in Gundlach et al. (2023) were presented throughout the experiment, while in our design, during the anticipatory period no visual stimuli were presented, which allowed differential anticipatory alpha power patterns to better manifest; it is known that visual stimulation suppresses alpha power and therefore may diminish the ability to detect different patterns of alpha power modulation across different attention conditions.

### Hierarchical organization of feature attention control

From the decoding accuracy time course, we observed significant <u>above-chance decoding</u> between attend-motion and attend-color conditions as early as ∼180 ms after cue onset. In addition, extending the MVPA analysis to classifying attended the specific feature attributes within a feature dimension, we found a delay in the significant above-chance decoding for attend-up versus attend-down and attend-blue versus attend-green, with significant classification not beginning until ∼385 ms. These onset latencies suggest a hierarchical model of feature attention control where the selection of the attended feature dimension precedes that of the attended feature attribute within a dimension.

Hierarchical control of feature attention has been proposed before based on the functional-anatomical patterns of activity in frontal-parietal cortex using fMRI. Liu and Hou (Liu & Hou, 2013) found that the spatial distribution of neural activity within the fronto-parietal cortex was more similar for within-feature (e.g., blue vs. green) and across feature dimensions (e.g., blue vs. upward motion) contrasts compared to between feature and spatial attention contrasts (e.g., blue vs. left visual field), which evoked more distinct neural patterns. That is, the functional-anatomical patterns suggested a hierarchical grouping based on task goals (spatial vs. feature attention). Such a functional-anatomical hierarchy could also give rise to a <u>temporal hierarchy</u>, as demonstrated in our EEG findings. Our results also align with the models of feature attentional selection in visual working memory (Brady et al., 2011; Hajonides et al., 2020; Töllner et al., 2015) and visual search experiments (Nako et al., 2014). According to the dimensional feature bundle model (Töllner et al., 2015), selecting targets based on visual features, such as a color or a shape, operates in three distinct levels, where the bottom level represents the attended feature attribute (e.g., blue for color or triangle for shape), the middle level is hypothesized to integrate the representations of the feature attributes at a dimensional level encompassing all the color or shape representations, and the top level integrates the representations of the respective feature dimensions. Niklaus et al. (2017), in a series of behavioral experiments, investigated the dimensional feature bundle model in the context of dimensional selection using a retroactive cueing design. These authors presented objects with varying colors or orientations (their Experiment 3), following a variable delay period, participants were presented with a retro-cue instructing participants to remember either the color or orientation of one of the targets. Following the retro-cue, on 50% of the trials, instead of presenting the probe for the response, participants were presented with a second retro-cue shifting their focus to either the color or orientation of a <u>different object</u> that was presented in the initial array. The switching costs from one stimulus to the other, measured using the difference in the error rates, were <u>minimal</u> if the new object (retro-cue 2) shared the <u>same</u> dimension as the one cued earlier (retro cue 1), in contrast to a condition were the participants were cued to report the characteristics of an un-cued dimension; e.g., if the first retro-cue required the participants to report the color of object 3 in the stimulus array and the second retro-cue required participants to identify the <u>orientation</u> (cross-dimensional switch) of object 1, then error rates were higher compared to the case when the second retro-cue asked participants to report the color of the object 1 (intra-dimensional switch). These findings support the presence of active representations of both the feature dimension and the attended feature attribute as opposed to just representing the specific feature attributes, because if the feature attributes alone were represented in the working memory as an independent unit, then the switching costs should have been identical.

Nako and colleagues (2014) investigated the role of category-based attentional guidance using visual search tasks by presenting alphanumeric arrays and instructing participants to search for one, two, or three targets (digits or letters) or targets defined based on categories (digits among letters, or vice versa). They also included catch trials (foil trials) where the search array did not contain the target item (e.g., A) but had an item within the same category (e.g., R). Analyzing the behavior and N2pc components, they report that both targets and foil trials evoked robust N2pc components, suggesting that in addition to arrays with target, arrays without the target but with category-matched items drew participants’ attention. Further, assessing the time course of N2pc components, they report the N2pc amplitudes between foil and target trials were comparable within 200 ms after the search array onset. However, from 200 ms after the search array onset, the target trials evoked stronger N2pc components, compared to the foil trials, suggesting that the stimulus processing in the initial stages is category-specific before progressing to item-specific attentional processing. Our proposed hierarchical control model of feature attention extends these models to the implementation of the top-down anticipatory attention control by demonstrating a temporal difference in the formation of dimensional-level attentional states and that of attribute-level attentional states.

Our findings can be described in terms of the Gating by Inhibition Model of attention control (Jensen & Mazaheri, 2010). In line with this model, our data show that top-down feature attention control occurs in two critical sequential stages involving alpha mechanisms. The first stage would involve suppression of the unattended feature dimension (increased alpha) and enhancement of the attended feature dimension (decreased alpha), giving rise to decodable differences in the alpha power topography. For our data, this stage corresponds to attention being directed to the color of the moving dots while ignoring the direction of motion or vice versa, and it onsets ∼180 ms after cue onset. In the second stage the top-down control selectively modulates the patterns sensitive to the attended attribute within the attended dimension while suppressing the unattended attribute within the same attended dimension, again leading to decodable differences in the alpha power topography the onset for this stage occurs around 385 ms after cue onset. While the Gating by Inhibition Model provides an interpretational framework for our findings, our results, in turn, can be seen to enrich the model by providing timing information on two stages of feature attention control.

### Relation with other relevant works in the literature

Our interpretation framework relies on the notion that increased amplitude of alpha power represents suppression of the activity in neural structures underlying ignored features(Jensen, 2024; Xia et al., 2024). For example, the dorsal stream is suppressed when color is attended, whereas the ventral stream is suppressed when attention is directed to motion (Snyder & Foxe, 2010; Verghese et al., 2013). Similar suggestions have been put forth in spatial attention studies (Fiebelkorn et al., 2018; Kelly et al., 2006; Sauseng et al., 2005; Wang et al., 2022; Worden et al., 2000; Wöstmann et al., 2019; Yang et al., 2023). Rihs et al., 2007, using a covert spatial attention task, cued participants to attend one of eight spatial locations organized around an imaginary sphere. Analyzing the post-cue alpha activity 250 ms before the onset of targets, they reported an increase in alpha power in the posterior electrodes *ipsilateral* to the attended location, indicating that alpha power serves to suppress the unattended dimension. Samaha et al. (2016) replicated this finding using an MVPA approach showing that classifiers trained using alpha activity during the cue-target interval were able to predict the attended location on a given trial.

Our findings also align with the feature similarity gain model of attention (Treue & Martínez Trujillo, 1999) which suggests that attention enhances the gain of the neurons based on the preference of the neurons for the attended feature. For example, during the attend blue condition, neurons tuned to color dimension and also to the blue stimulus attribute are enhanced while the neurons preferring the opposite motion dimension are suppressed; the opposite is true when attention is directed to up or down moving dots where the color dimension is suppressed (Chen et al., 2012). We propose that the feature gain modulation is implemented in two stages where the gain of the attended and unattended feature dimension is regulated first before narrowing the attention-related biasing to the attended feature attribute.

## Conclusions

Using EEG recordings of oscillatory activity in the alpha band, and multivariate decoding analyses, we show that top-down voluntary feature attention control is implemented in the brain in a hierarchical fashion. The more general feature category (feature *dimension*: color vs. motion) of the to-be-attended stimuli is biased *before* the biasing of more specific categories (feature *attributes*: blue vs. green; upward motion vs. downward). These results support a hierarchical attention control model for feature attention in humans.

## Acknowledgements

This work was supported by National Institutes of Health grant MH117991 and National Science Foundation grant BCS-2318886 to G.R.M. and M.D. We thank Sai Katta for help with data collection, and our colleagues in the Center for Mind and Brain for helpful discussions.

## Author Contribution

**Sreenivasan Meyyappan (lead):** Conceptualization; formal analysis; methodology; writing. **Mingzhou Ding** and **George R. Mangun**: Conceptualization; funding acquisition; writing.

